## Supplementary figures and legend and table for "MYC-ATF4-ASS1 axis governs intracellular arginine synthesis and dictates the immune microenvironment in melanoma"

**Supplementary Figure 1. A combination therapy of BRAF/MEK inhibitors and ADI-PEG20 reduces organoid proliferation.** WCM214 (a) or WM4631 (b) Organoids from PDX were treated with DMSO, D/T, ADI, or D/T+ADI for 7 days. CellTiterGlo assay were performed to examine the organoid proliferation. * indicate $p<0.05$.

**Supplementary Figure 2. ASS1 expression in melanoma patients or PDX.** (**a**) Ass1 expression is lower in melanoma patients compared to normal tissue from TCGA dataset. (**b** and **c**) The expression profile of ASS1 in PDX datasets of Dr. Herlyn's lab. Two batches both show most of the PDX express low level of ASS1.

**Supplementary Figure 3. MYC ChIP-seq and its targets.** (**a**) WM4380-2 MYC Wiltype (WT) or knockout (KO) cells were subjected to ChIP-seq to identify the direct targets of MYC. Diffential peaks were annotated and shown as volcano plot. (**b**) ARHGDIA, as a positive control of MYC direct target, was pulled down in the MYC-WT cells but not in the MYC-KO cells, demonstrating the reliability of the pulled-down targets of MYC-ChIP-seq.

**Supplementary Figure 4. The annotation of cell types from single cell RNA sequencing of Ass1-WT or Ass1-KO melanoma.** *Cspg4* is a melanoma marker, *Itgam* is a macrophage marker, *Vwf* is a endothelial cell marker, *Ptprc* is a total immune cell marker, *Cd3e* is a total T cell marker, while *Foxp3* is regulatory T cell marker.

**Supplementary Figure 5. Global reduction of Cell-Cell-Communication in the Ass1-KO melanoma compared to the wildtype melanoma from single cell analysis.**

### Sup Fig. 1

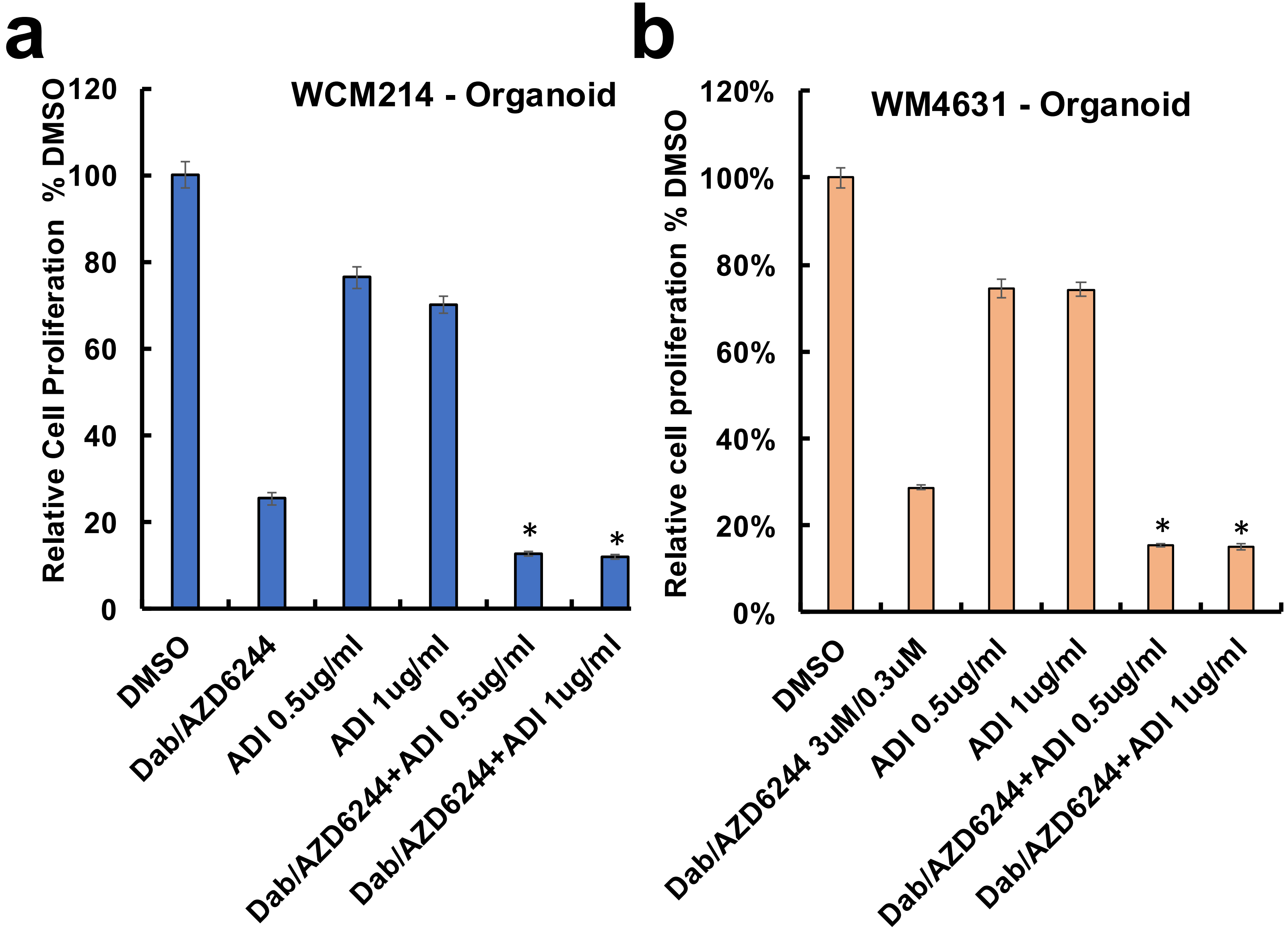
a
WCM214 - Organoid
Relative Cell Proliferation % DMSO
120
100
80
60
40
20
0
*
*
DMSO
Dab/AZD6244
ADI 0.5ug/ml
ADI 1ug/ml
Dab/AZD6244+ADI 0.5ug/ml
Dab/AZD6244+ADI 1ug/ml
b
WM4631 - Organoid
Relative cell proliferation % DMSO
120%
100%
80%
60%
40%
20%
0%
*
*
DMSO
Dab/AZD6244 3uM/0.3uM
ADI 0.5ug/ml
ADI 1ug/ml
Dab/AZD6244+ADI 0.5ug/ml
Dab/AZD6244+ADI 1ug/ml

### Sup Fig. 2

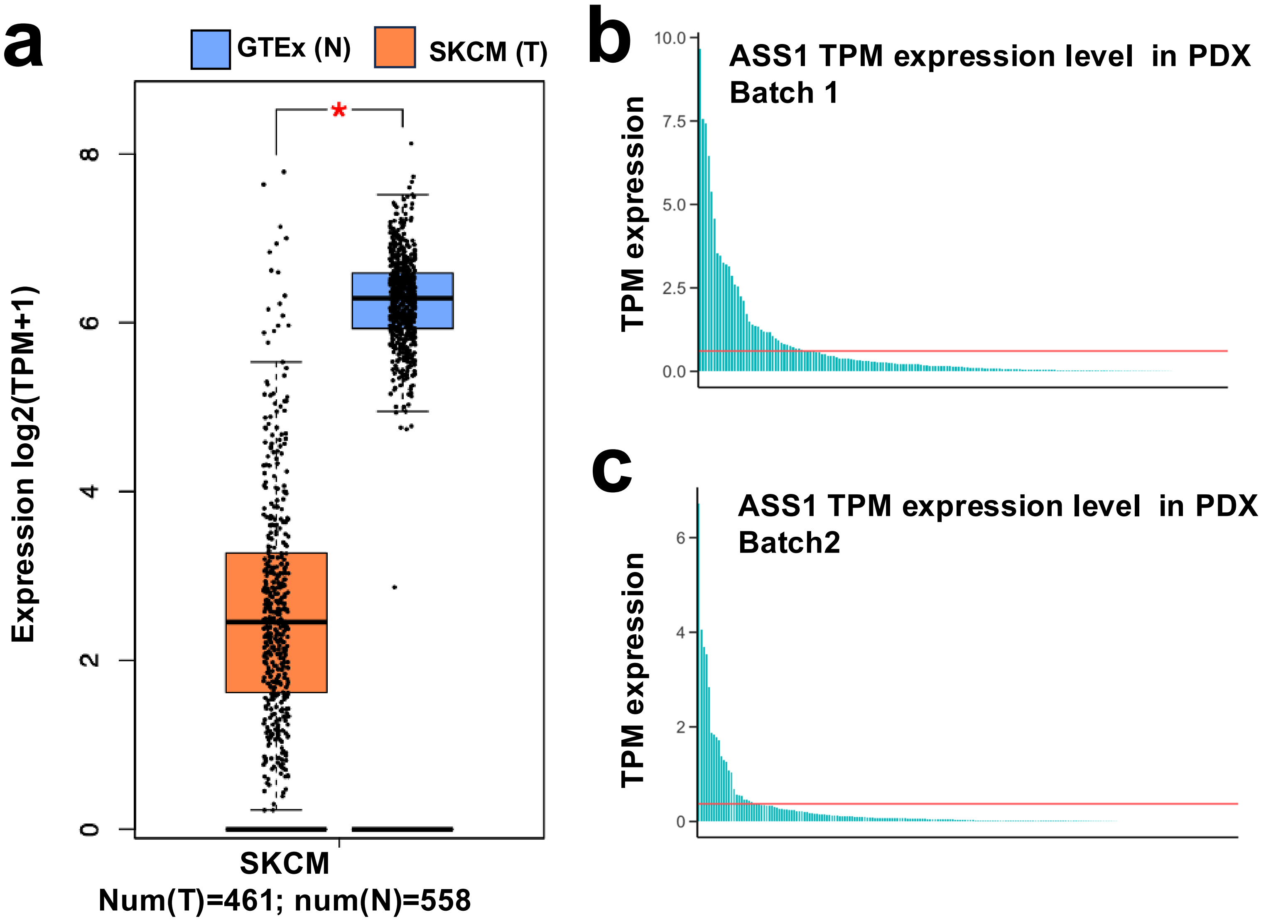
a
GTEx (N)
SKCM (T)
*
Expression log2(TPM+1)
0
2
4
6
8
SKCM
Num(T)=461; num(N)=558
b
ASS1 TPM expression level in PDX Batch 1
TPM expression
0.0
2.5
5.0
7.5
10.0
c
ASS1 TPM expression level in PDX Batch2
TPM expression
0
2
4
6

### Sup Figure 3

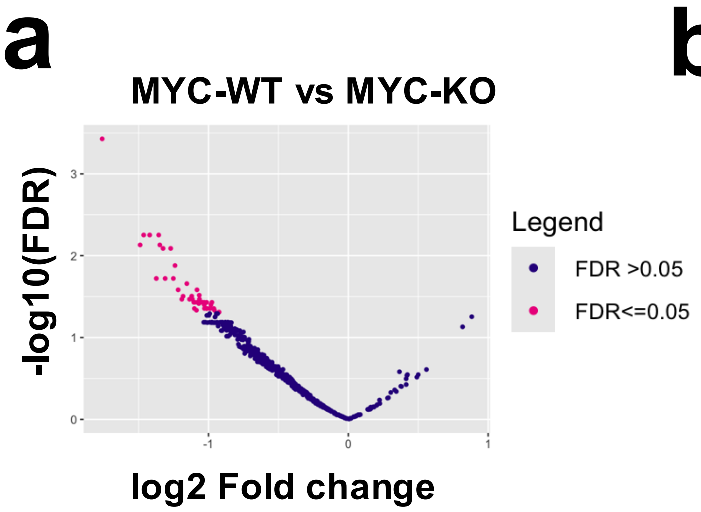
a
MYC-WT vs MYC-KO
-log10(FDR)
log2 Fold change
Legend
FDR >0.05
FDR<=0.05

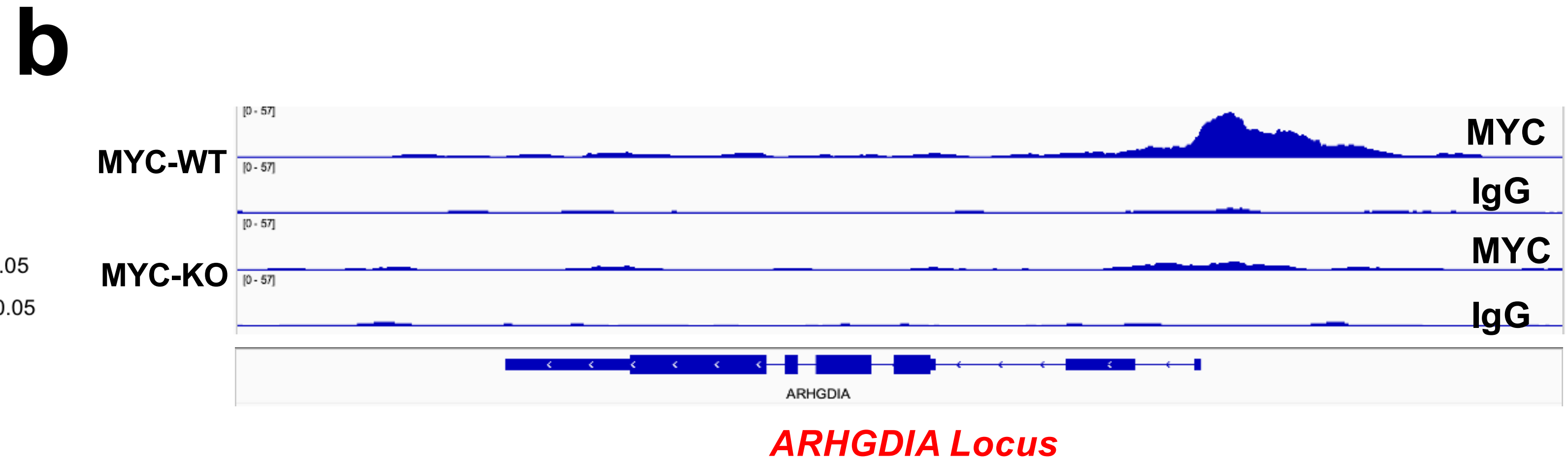
b
MYC-WT
MYC-KO
MYC
IgG
MYC
IgG
ARHGDIA
ARHGDIA Locus

### Sup Figure 4

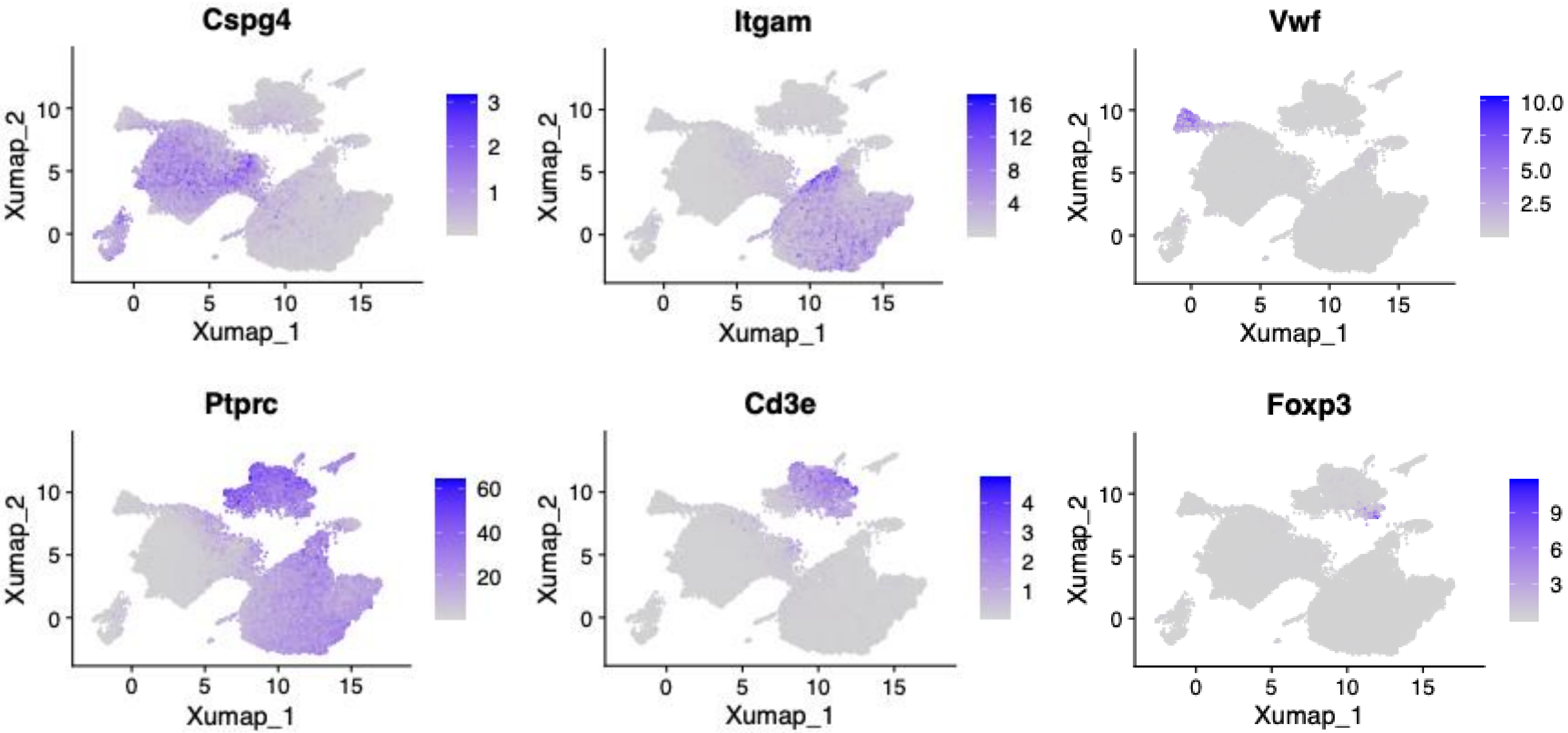

### Sup Figure 5

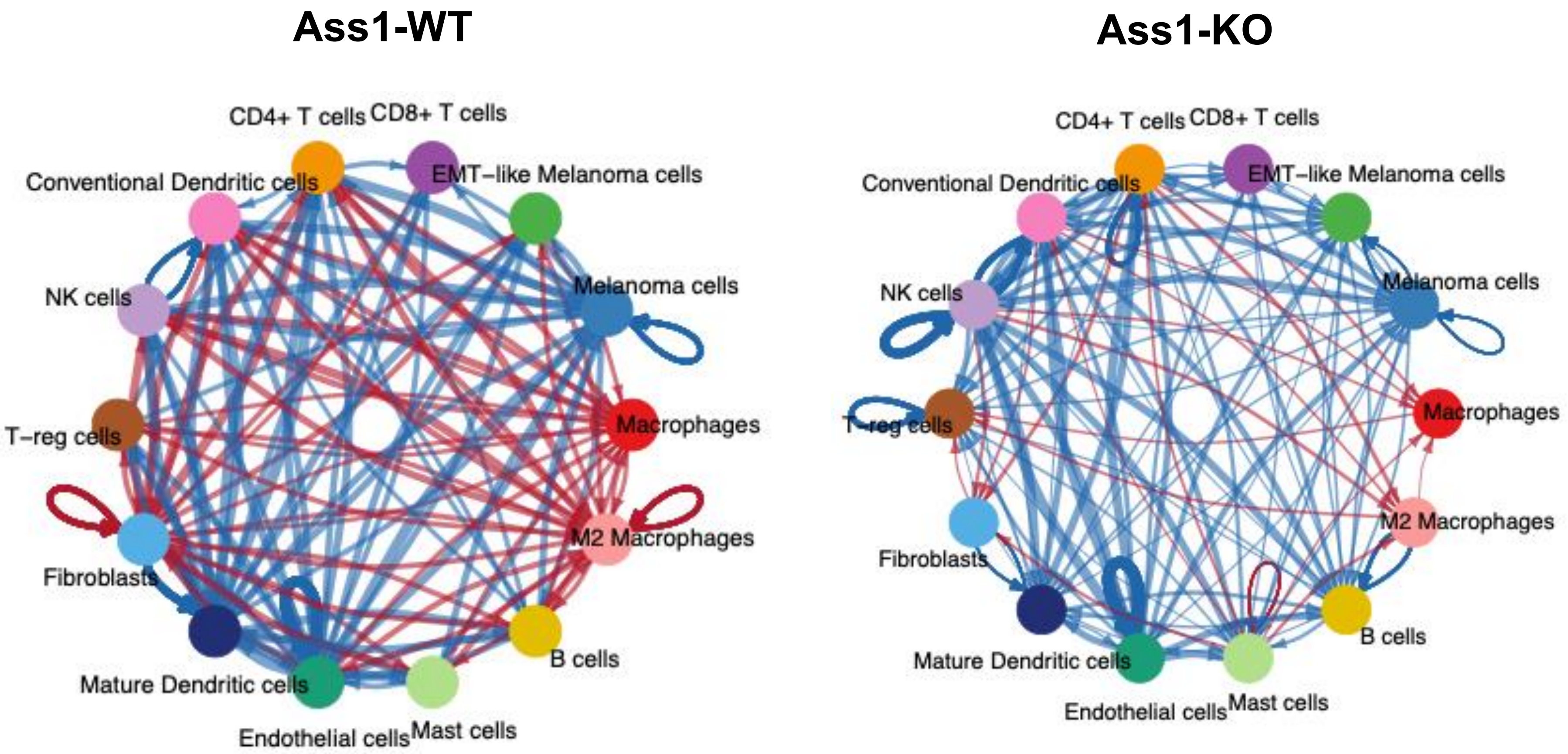


Differential number of interactions

**WCM214**

Methods: Data from 41 days to 71 days were used in statistical analysis with mixed-effect model. The tumor volume at 41 days was used as the baseline data at pre-treatment.

Table 1. Linear mixed-effects model estimated tumor growth rates by group

| group | estimated tu | Std. Err. | z | P>z | [95% Conf. | Interval] |
|---|---|---|---|---|---|---|
| Control | 100.7608 | 18.36808 | 5.49 | 0 | 64.75999 | 136.7616 |
| D/T | 43.48338 | 3.251713 | 13.37 | 0 | 37.11014 | 49.85663 |
| ADI | 19.82882 | 2.637404 | 7.52 | 0 | 14.6596 | 24.99804 |
| D/T+ADI | -8.282949 | 1.816575 | -4.56 | 0 | -11.84337 | -4.722527 |

Table 2. P-values from mixed-effects model comparing the estimated tumor growth rate between groups

| **group** | **Control** | **D/T** | **ADI** |
|---|---|---|---|
| **Control** | | | |
| **D/T** | **<0.0001** | | |
| **ADI** | **<0.0001** | **0.046** | |
| **D/T+ADI** | **<0.0001** | **<0.0001** | **0.0295** |

**WM4631**

Methods: Data from 16 days to 43 days were used in statistical analysis with mixed-effect model. The tumor volume at 16 days was used as the baseline data at pre-treatment.

Table 3. Linear mixed-effects model estimated tumor growth rates by group

| group | estimated tu | Std. Err. | z | P>z | [95% Conf. | Interval] |
|---|---|---|---|---|---|---|
| Control | 56.34228 | 8.271578 | 6.81 | 0 | 40.13029 | 72.55428 |
| D/T | 12.70395 | 5.438563 | 2.34 | 0.019 | 2.044563 | 23.36334 |
| ADI | 24.67513 | 3.217337 | 7.67 | 0 | 18.36927 | 30.981 |
| D/T+ADI | 0.0652582 | 1.295674 | 0.05 | 0.96 | -2.474217 | 2.604733 |

Table 4. P-values from mixed-effects model comparing the estimated tumor growth rate between groups

| **group** | **Control** | **D/T** | **ADI** |
|---|---|---|---|
| **Control** | | | |
| **D/T** | **<0.0001** | | |
| **ADI** | **0.0001** | **0.1291** | |
| **D/T+ADI** | **<0.0001** | **0.0979** | **0.0016** |
